# Metalign: Efficient alignment-based metagenomic profiling via containment min hash

**DOI:** 10.1101/2020.01.17.910521

**Authors:** Nathan LaPierre, Mohammed Alser, Eleazar Eskin, David Koslicki, Serghei Mangul

## Abstract

Whole-genome shotgun sequencing enables the analysis of microbial communities in unprecedented detail, with major implications in medicine and ecology. Predicting the presence and relative abundances of microbes in a sample, known as “metagenomic profiling”, is a critical first step in microbiome analysis. Existing profiling methods have been shown to suffer from poor false positive or false negative rates, while alignment-based approaches are often considered accurate but computationally infeasible. Here we present a novel method, Metalign, that addresses these concerns by performing efficient alignment-based metagenomic profiling. We use a containment min hash approach to reduce the reference database size dramatically before alignment and a method to estimate organism relative abundances in the sample by resolving reads aligned to multiple genomes. We show that Metalign achieves significantly improved results over existing methods on simulated datasets from a large benchmarking study, CAMI, and performs well on *in vitro* mock community data and environmental data from the Tara Oceans project. Metalign is freely available at https://github.com/nlapier2/Metalign, along with the results and plots used in this paper, and a docker image is also available at https://hub.docker.com/repository/docker/nlapier2/metalign.

## Introduction

Microorganisms are ubiquitous in almost every natural setting, including soil ^1^, ocean water ^2^, and the human body ^3^, and they play critical roles in the functioning of each of these systems ^4, 5^. Traditional culture-based analysis of these microbes is confounded by the well-known fact that many microorganisms are not culturable in standard laboratory settings ^4, 6^. Analysis of lab-cultured organisms also cannot capture the complex community dynamics in real microbial ecosystems ^4^. Powered by modern high-throughput sequencing, the field of metagenomics, or the analysis of whole microbial genomes recovered directly from their host environment, is vital to understanding microbial communities and their functions ^4, 5^. Predicting the presence and relative abundance of organisms in a metagenomic sample (referred to as “profiling”) is one of the primary means of analyzing a metagenomic sample ^7, 8^. In comparison to assembly, profiling has the advantages of being computationally simpler and more effective at identifying low-abundance organisms ^8^.

A recent widely-cited comprehensive benchmarking study, the Critical Assessment of Metagenome Interpretation (CAMI) ^7^, evaluated ten widely-used profiling methods on a variety of simulated metagenomic datasets. Notably, the two best profiling methods in precision (i.e., false positive rate) were the two methods ranked lowest for recall (i.e., false negative rate); similarly, the two best methods for recall were two of the three methods ranked lowest for precision ^7^. This makes sense intuitively as it is easy to select only a few high-confidence species as being present or to claim almost all known species are present; balancing precision and recall presents a substantial challenge. Nevertheless, this discrepancy presents a difficult trade-off to researchers, who must choose between failing to identify most organisms or falsely identifying many organisms. Similarly, the top-ranked method in L1-error and Weighted UniFrac ^9, 10^, measures of error in abundance estimation and taxonomic composition, respectively, had the second-worst precision ^7^, indicating another difficult trade-off. Methods that were ranked among the worst in recall and abundance estimation in the CAMI competition ^7^ have been used in large-scale efforts to analyze human^11^ and city metro ^12^ microbiomes and downstream analyses linking the microbiome to host genetics ^13^ and diseases such as colorectal cancer, potentially impacting the findings of these important studies.

In order to address common obstacles to metagenomic analyses, we developed Metalign, an efficient alignment-based metagenomic profiling method that achieves a strong balance of precision and recall with runtimes comparable to state-of-the-art methods. Alignment-based profiling is regarded as highly accurate, but aligning millions of reads against a reference database of tens to hundreds of gigabytes (GB) in size is computationally infeasible. To avoid this issue, Metalign employs a high-speed, high-recall pre-filtering method based on the mathematical concept of Containment Min Hash ^14^, which identifies a small number of candidate organisms that are potentially in the sample and creates a subset database consisting of these organisms. Empirically, this approach reduced our comprehensive NCBI-based database of 787 GB more than 100-fold, with some variance depending on the diversity of the sample. We limit false positives by performing a highly accurate alignment step on the subset database, which both handles the reads that align uniquely to one genome and the reads that align to multiple genomes. Metalign then profiles the organisms that reach a certain threshold amount of reads uniquely aligned to their genome, along with other optional metrics (see Methods).

We show that Metalign significantly outperforms the ten state-of-the-art profiling methods that were evaluated in the CAMI competition ^7^ and achieves strong results on *in vitro* mock community data ^15^. We demonstrate that Metalign identifies a diverse, well-supported set of microbial clades, with competitive runtime, in real environmental data from the Tara Oceans project ^16^. Metalign should benefit researchers seeking to efficiently obtain highly accurate metagenomic community profiles, enabling more accurate scientific discovery and downstream analysis.

## Results

### Methods overview

Aligning millions of reads to reference databases — which are hundreds of GB in size — is computationally infeasible. However, with an effective pre-filtering stage (**Figure 1a**), highly accurate alignment can be performed on the small pre-filtered database. We use KMC ^17^ to generate k-mers for the reads and each reference genome, and then we utilize an implementation of the theoretical concept of Containment Min Hash ^14^ to estimate the percent of k-mers in each reference genome that are also present in the reads (Methods). Intuitively, this gives us an estimate of how likely it is for each reference genome to be present in the sample. We place in a subset database only the reference genomes above a certain percentage cutoff threshold, to which we align the reads using the fast modern aligner Minimap2 ^19^. Finally, we estimate the relative abundances of microbes in the sample by combining information from reads that map uniquely to one genome with those that align to multiple genomes (see Methods).

**Figure 1.**
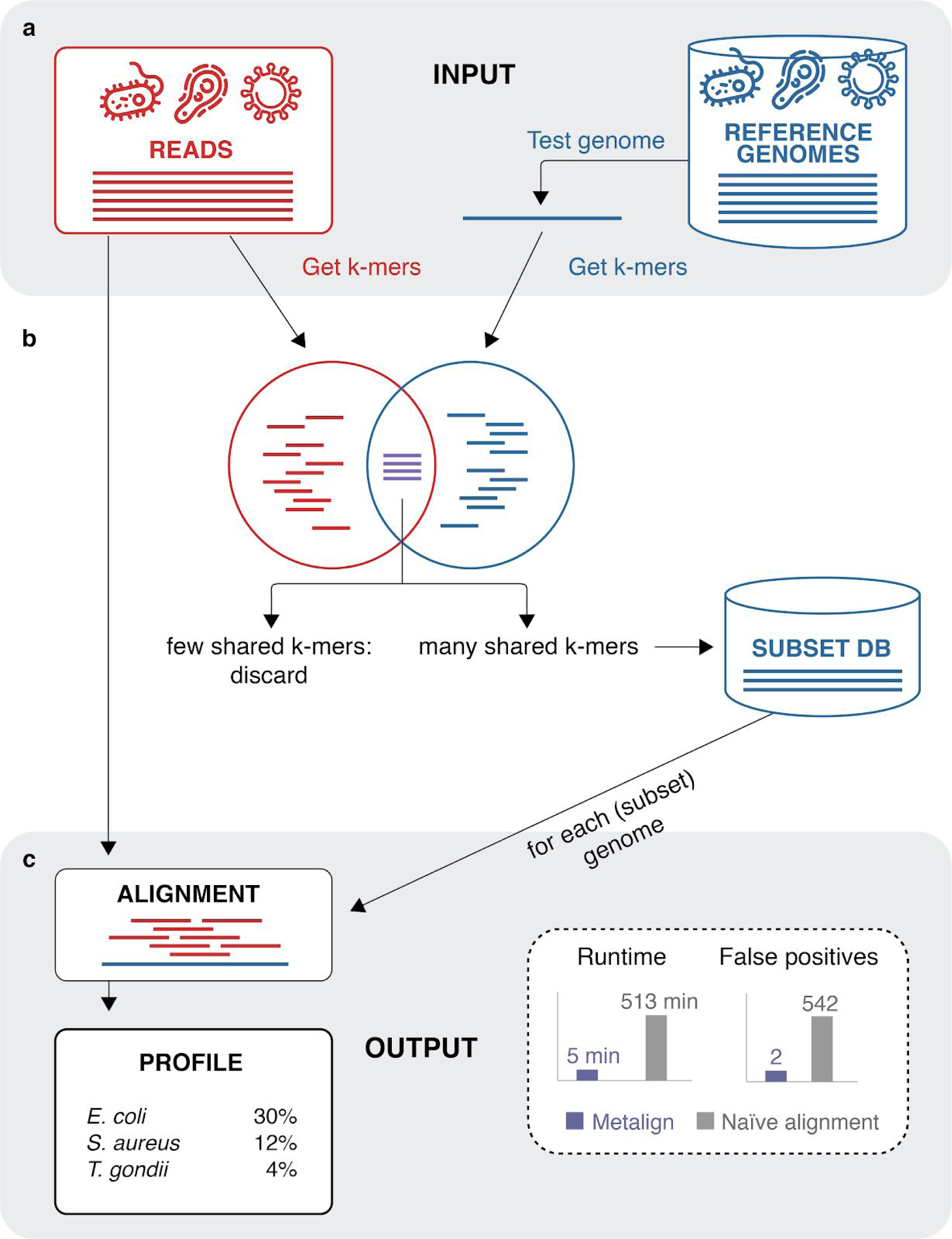
Metalign overview. Panel A shows the input to our method, sequencing reads and our reference database, and the k-mer counting step using KMC ^17^. The pre-filtering stage (Panel B), based on an implementation of the theoretical concept of Containment Min Hash ^14^, quickly estimates the percentage of k-mers in each reference genome that are also in the reads. We then select a small “subset” database consisting of reference genomes above a certain percentage threshold. As shown in Panel C, we then perform alignment between the reads and the reference genomes in the subset database, outputting a profile in the standardized, community driven format used by OPAL ^18^ and CAMI ^7^. We found that, when applying Metalign to *in vitro* mock community data compared with naive alignment without pre-filtering, we reduced runtime from 513 minutes to 5 minutes, and false positive genera from 542 to 2.

We tested the efficacy of our CMash database prefiltering step (Methods) by running alignment with Minimap2 on our full unfiltered database and on the reduced database produced by our pre-filtering step. We then applied the alignment and profiling stage of Metalign. We performed these experiments on a mock community dataset consisting of 300,969 reads, obtained from Peabody et al. ^15^ Alignment and profiling with the full database took approximately 513 minutes, while alignment and profiling with the CMash-reduced database—enabled by Metalign—took approximately 5 minutes. Additionally, the pre-filtering step reduced noise in the form of spurious alignments to organisms that were not present in the data. Consequently, while both strategies detected all genera present in the data, profiling using the full unfiltered database produced 542 false positive genera, while profiling using the pre-filtered database produced only two false positives. This experiment highlights the significant improvements in speed and precision enabled by Metalign’s CMash-based pre-filtering step.

### Metalign significantly outperforms existing methods on CAMI simulated data

The Critical Assessment of Metagenome Interpretation (CAMI) ^7^ provides the most comprehensive and in-depth evaluation of metagenomic profiling, binning, and assembly methods to date. In the profiling competition, many of the most well-known methods were evaluated on a variety of simulated datasets that modeled real-life challenges, such as various community diversities and confounding sequences from high-abundance plasmids and novel viral strains. We evaluated the performance of the top-ranked methods in terms of several metrics: recall, precision, F1 score, Jaccard index, L1 norm error, and Weighted UniFrac (Supplementary information). In total, there were eight datasets: one low-diversity community, two medium-diversity communities, and five high-diversity communities. Each dataset consisted of 15 Gbp of sequence data. (Further details on these communities are available in the CAMI paper ^7^.)

We ran Metalign on these eight datasets and compared our results with publicly-released results from state-of-the-art existing methods using the CAMI-affiliated evaluation software OPAL ^18^ (**Figure 2**). We only included methods that had submitted results for all eight datasets. Because Taxy-Pro ^20^ had two submissions, for brevity we included only the second one (“Taxy-Pro2”) Taxy-Pro1 did not stand out in any particular performance metric, and Taxy-Pro2 had the best Weighted UniFrac (**Figure 2b**).

**Figure 2.**
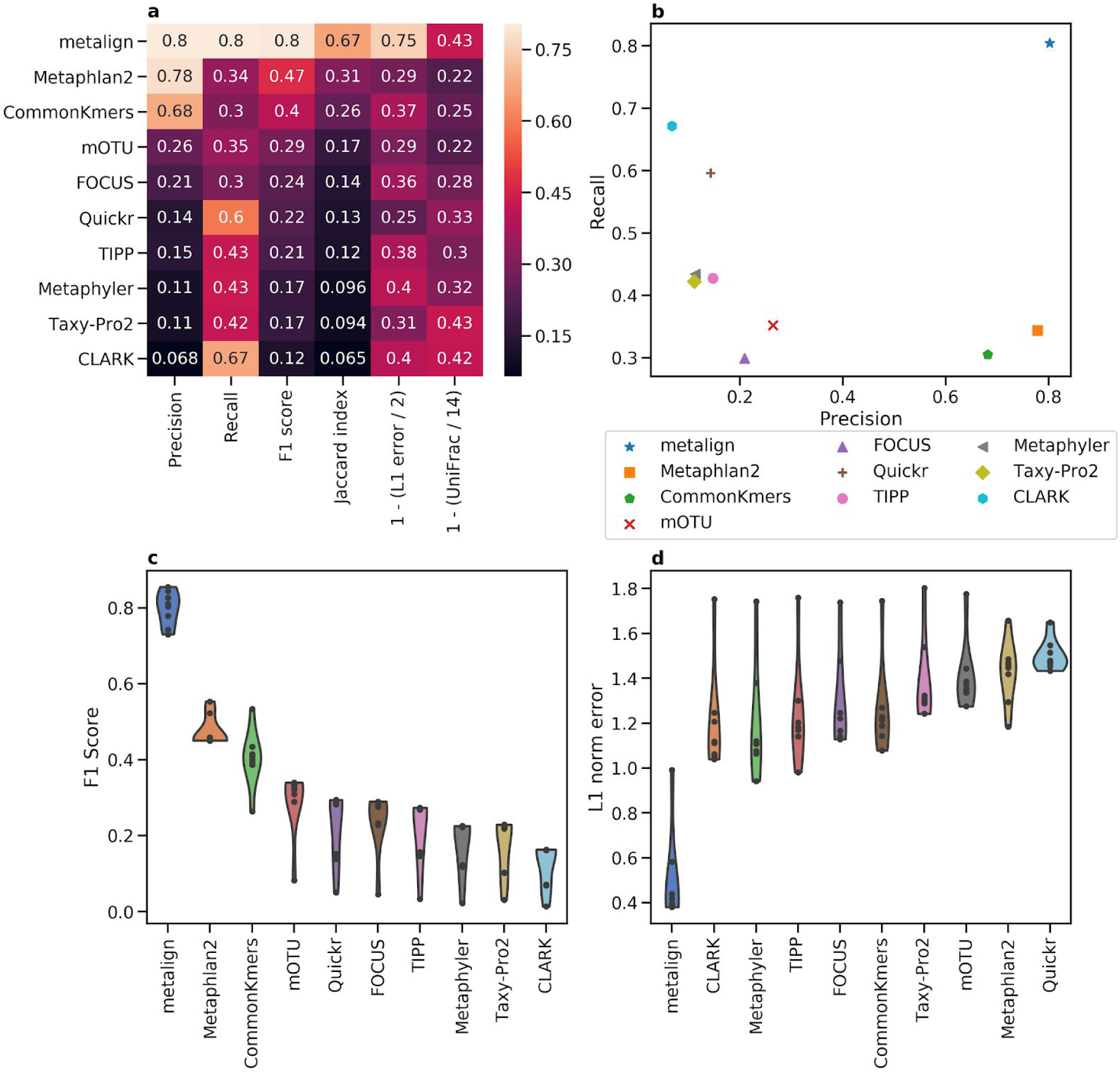
Comparison between Metalign and methods submitted to the first CAMI competition on the genus level. (A) Heatmap showing Precision, Recall, F1 score, Jaccard index, and 0-to-1 scaled versions of L1 error and Weighted UniFrac, such that higher is better for each column, averaged across all datasets. (B) A scatter plot of Precision (x-axis) versus Recall (y-axis), averaged across all datasets. (C and D) Violin plots of F1 score (C; higher is better) and L1 error (D; lower is better), respectively, where the x-axis contains the methods, the y-axis represents the metric, and each dot represents one dataset.

Averaging across all datasets on the genus level, five of the ten methods fail to achieve either 50% precision or 50% recall, and only Metalign was above 50% in both of these metrics—achieving roughly 80% in both precision and recall (**Figure 2a-b**). Similarly, Metalign outperformed, in both the F1 Score and Jaccard index, all other methods on the genus level by more than 30%. Further, Metalign had the best L1 error by about 58.09% (**Figure 2a-b**). Metalign was consistent across all datasets of varying diversity, outperforming other methods on the genus level in F1 Score and L1 norm error in all eight datasets **(Figure 2c-d)**. Metalign also outperforms other methods on the species level, except in precision where it is second-best after MetaPhlAn2; however, its recall is about five times as high as that of MetaPhlAn2 on the species level (**Figure S1**). Averaging across all taxonomic levels, Metalign outperforms all other methods in recall, F1 score, Jaccard index, and L1 norm error, while finishing third in precision (**Figure S2**). Metalign outperformed the next best method in F1 Score and Jaccard index by more than 10% (**Figure S2a**). On the rank-independent metric Weighted UniFrac, Metalign finishes second among the methods, nearly the same as the top Method, Taxy-Pro (**Figure 2a**).

Overall, Metalign outperformed other methods in abundance estimation and presence/absence balance across a variety of datasets and taxonomic ranks. In terms of CPU time, Metalign took between 38 and 67 minutes per sample, with higher diversity samples taking longer; timing results for other methods were not available.

Methods submitted to the first CAMI competition were allowed to use any reference database. However, the number of reference genomes provided by NCBI has grown since the CAMI competition took place in 2017. This relative abundance in reference genomes allows Metalign to use a more comprehensive database than prior methods had access to. However, re-running MetaPhlan2 using its most up-to-date reference database yielded similar results to its results in the original competition. Additionally, most of the other methods above have high recall and low precision, and more comprehensive databases would likely only exacerbate that discrepancy.

### Metalign outperforms other methods across cutoff settings on mock community data

While simulated data often fails to fully capture the noise present in environmental microbial communities, a ground truth is not available for the latter, making the calibration and comparative evaluation of competing methods difficult^21^. *In vitro* mock communities are specifically designed communities of microbes cultured in the laboratory which can then be sequenced. Such mock communities offer the benefit of being “real” data—albeit not from a natural environment—and having available ground truth.

One such dataset consists of 11 bacterial species and is available from a metagenomics benchmarking study performed by Peabody et al. ^15^. We evaluated Metalign against MetaPhlAn2 ^22^ and two new versions of popular methods from the CAMI ^7^ study: Kraken2 ^23^, with its abundance estimation companion method Bracken ^24^, and mOTUs2 ^25^. For all methods, we used their full standard databases and default settings. We evaluated presence/absence prediction, as ground truth relative abundance information is not available.

The Kraken family of methods is well-known to generate a number of very low abundance false positives ^7, 15^; therefore, it is common to employ some sort of abundance cutoff with their method ^15^. We evaluated the genus-level F1 score for these methods at a variety of cutoff thresholds (**Figure 3**). We excluded results reported as 0.0, including those with any number of trailing 0s.

**Figure 3.**
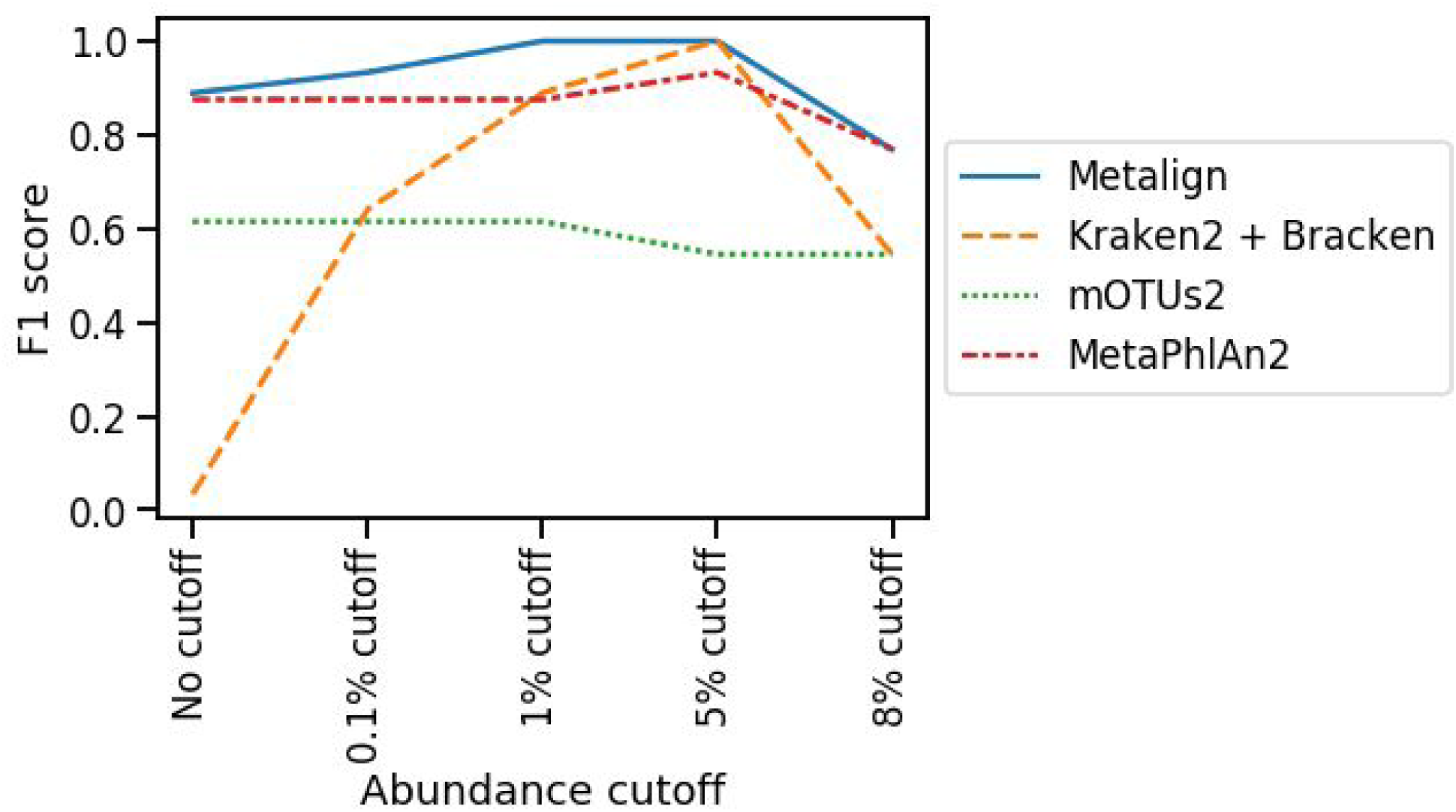
Genus-level F1 score at different abundance cutoff thresholds using *in vitro* mock community from Peabody et al. 2015 ^15^. For a given cutoff threshold, we calculated the F1 score while leaving out predicted genera below the threshold.

Metalign achieves the highest or tied-for-highest F1 score among the tested methods at all cutoff thresholds, having only two low-abundance false positives and no false negatives. MetaPhlAn2 performs almost as well, which we expected given a low-diversity community composed of common, high-abundance bacteria. Despite the low complexity of the sample, mOTUs2 did not predict most taxa down to the genus level and demonstrated poor performance in both recall and F1 score. Kraken2 + Bracken, as expected, generated numerous low abundance false positives. It is able to achieve a perfect F1-score with a cutoff threshold between about 1.8% and 7.1%, but users would not know this information in a real setting and may choose a different, less effective cutoff threshold. Metalign’s performance is strong across all thresholds, obviating the need for the user to set an arbitrary cutoff value.

### Metalign detects greater diversity of microbial communities in real world data

Mock communities are usually composed of species that have been previously cultured and cannot truly capture the amount of noise in an environmental sample. In addition, the mock community explored in our cutoff setting test is comprised of a small number of well-known bacterial species, and does not test the ability of methods to detect lower-abundance, higher-diversity, or less well-known clades. Tara Oceans, the largest Ocean microbiome sequencing project, provides the opportunity to address these concerns.

We tested Metalign, MetaPhlAn2 ^22^, Kraken2 ^23^ with Bracken ^24^, and mOTUs2 ^25^ on five prokaryote-isolated samples (**Supplementary Information**) from a deep chlorophyll maximum (DCM) layer of the ocean (**Figure 4**). The reads originally came in separate gzipped paired end files, which we decompressed and then interleaved with BBMap ^26^ before running the methods on this data. Results were then visualized with Krona ^27^ (**Figure 4a**, **Supplementary Figures S4-S7**).

**Figure 4.**
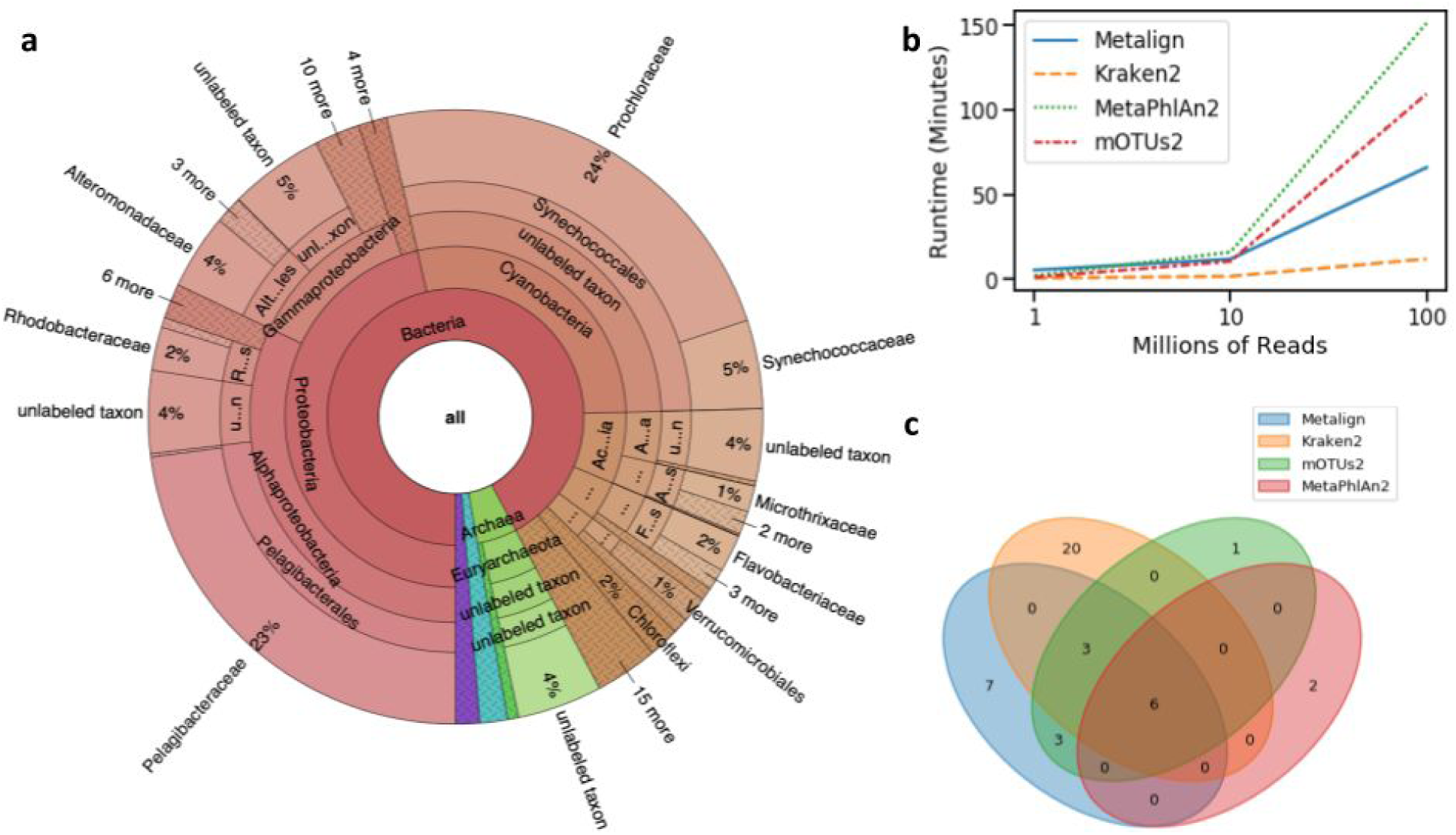
Comparison of Metalign’s performance to other methods on five deep chlorophyll maximum (DCM) layer samples from the Tara Oceans data. (A) Krona plot of the taxa found by Metalign, down to the family level, with abundances averaged over the five datasets. (B) Runtime scaling comparison between Metalign, Kraken2, MetaPhlAn2, and mOTUs2 on subsets of 1 million, 10 million, and 100 million reads from a large Tara Oceans sample. (C) Venn diagram based off of the phyla found by each method in the DCM samples, excluding phyla with <0.01% abundance.

Results from all methods agreed that the taxonomic families Pelagibacteraceae and Prochloraceae were highly abundant and that phyla Proteobacteria and Cyanobacteria dominated the samples. However, Metalign generally identified a more diverse set of high-level clades than did other methods. For example, MetaPhlAn2 did not identify the phylum Chloroflexi, whereas Metalign and Kraken2 correctly identified its presence (the latter with only 0.028% abundance, making it a candidate for filtering via abundance cutoff). Notably, Chloroflexi was found in the original Tara Oceans study ^16^ and in a subsequent comprehensive assembly analysis ^28^. Kraken2 did not identify the phylum Marinimicrobia, whereas Metalign, MetaPhlAn2, and the assembly-based Tara Oceans analysis did ^28^. Metalign identified more unique phyla than MetaPhlan2 or mOTUs2 did (**Figure 4c**). While Kraken2 identified the most unique phyla, many were low abundance, and the Kraken family of methods are prone to finding many low abundance false positives ^7, 15^. Ground truth for the data used here is not available, but Metalign’s results seemed largely in line with other, more comprehensive analyses of the data.

Kraken2 was able to achieve the fastest runtime, followed by Metalign, mOTUs2, and MetaPhlan2, respectively. All methods were able to evaluate even the largest individual sample (about 206 million reads) in fewer than six hours. To show the scalability of these methods to various sample sizes, we downsampled the largest sample to 100 million reads, 10 million reads, and 1 million reads, and we ran all methods on these samples (**Figure 4b**). All methods were capable of quickly processing 1 million and 10 million reads, but Metalign scaled better to 100 million reads than did MetaPhlAn2 or mOTUs2, while Kraken2 scaled the best. Metalign is therefore able to leverage an alignment-based approach while remaining competitive in runtime with current state-of-the-art methods.

## Discussion

We have developed a novel computational approach (Metalign) that is able to achieve a strong balance between precision and recall across a variety of datasets, community diversities, and taxonomic levels. This also obviates the need for the user to set an arbitrary abundance cutoff threshold for their data that could potentially eliminate true positive results. However, there is an inherent trade-off between precision and recall, and users must have the ability to modulate this trade-off based on their analytical needs. In addition to the option of applying a post-profiling abundance threshold, we allow users to modify the threshold used in our pre-filtering step, followed by standard alignment and profiling. We believe that the latter strategy is somewhat less arbitrary, as there is not necessarily a linear relationship between abundance and likelihood of presence. The latter approach allows the profiling step to adjust to a different set of organisms.

Several directions hold promise for further improvements to Metalign. The default containment index cutoff for inclusion of organisms in the reduced database is currently set in an empirical and somewhat arbitrary fashion, but preliminary theoretical results indicate that the containment index is related to the well-known Average Nucleotide Identity, which facilitates less arbitrary cutoff. The resolution of multi-aligned reads could be performed in a more robust manner at the expense of additional computational time; for example, by using base-level metrics such as quality scores and CIGAR strings ^29^. The standard of evidence used to determine the presence of an organism by Metalign could potentially be automatically modulated based on characteristics of the sample such as sequencing depth and estimated alpha diversity.

## Methods

### Database construction

In order to compile as comprehensive a reference database as possible, we used all NCBI microbial genome assemblies, including both complete and incomplete assemblies from both RefSeq and GenBank. The final database, as of the writing of this paper, consisted of 199,835 organisms totaling 787 GB in size. We intend to continue updating it alongside NCBI database updates. We use rsync to download all NCBI genomes and then filter out unwanted taxa such as animals and plants. The script for this procedure is available on our GitHub (https://github.com/nlapier2/Metalign).

### Database pre-filtering with CMash

Aligning millions of reads to reference databases of hundreds of GB in size is computationally infeasible. However, we show in this paper that aligning to a much smaller pre-filtered database can be done in a reasonable amount of time. Our pre-filtering stage uses a k-mer based approach to focus on high speed and high recall (low false negative rate). First, KMC ^17^ is used to enumerate the k-mers in the reads, with the k-mers of the reference genomes having been pre-computed by KMC, and then intersect these sets. We then utilize the containment MinHash similarity metric (presented theoretically in ^14^ via an implementation by one of the coauthors (Koslicki) called CMash to efficiently estimate the similarity/containment index) between each reference genome and the input sample. The containment index is closely related to the Jaccard index, and, in this case, refers to the percent of k-mers in a reference genome that are also present in the reads.

Given the estimates for the containment index for each reference genome, we select all reference genomes above a certain cutoff threshold for inclusion in a new, reduced database on which to perform alignment. We currently set the cutoff to

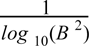

where *B* is the size of the file in bytes, as this value empirically gave us a good balance between recall and precision in our downstream results. This cutoff threshold is designed to scale inversely with the size of the reads file; the greater an imbalance between the number of k-mers in a reference genome and the number of k-mers in the reads, the greater the bias towards zero will be in the estimation of the containment index.

We allow users to modulate the cutoff threshold to suit their needs, as it controls the balance between precision and recall. In particular, we include a pre-built --sensitive option in our software that sets the cutoff threshold to zero, meaning that if CMash estimates the containment index for a genome to be nonzero, the genome will be included in the reduced database. By default, we keep only one strain per species (the one with the highest containment index) above the cutoff threshold, and we discard extra strains. Metalign is capable of strain-level profiling via an optional software flag; however, in practice, profiling strains tends to slow down computation. Based on these user-set options, we construct a reduced database on which to perform alignment and profiling.

### Alignment and profile generation

Alignment of the reads to the pre-filtered database is performed using Minimap2 ^19^, which we empirically observed to generate similarly accurate results to older alignment methods but in much less time. The main challenges with profiling given mapping results are in handling multi-aligned reads (reads successfully aligned to multiple reference genomes) and how to determine what amount of evidence is needed to consider a species present. These choices affect trade-offs between abundance estimation, precision, and recall.

Multi-aligned reads are resolved according to the uniquely-mapped abundances of the organisms that a read is aligned to. First, we calculate the abundance estimates using only the uniquely-aligned reads, holding aside reads that align to multiple genomes. Next, for each read that aligns to multiple genomes, we assign its abundance to the genomes it aligns to proportionally to the number of reads that align uniquely to each of those genomes. As a concrete example, assume species A has 6000 uniquely-aligned reads, and species B has 2000. A read aligned to both species A and species B thus has 75% of its bases assigned to species A, and 25% of its bases assigned to species B. Note that this means that we do not perform explicit read classification/binning; rather, we try only to obtain the relative abundances of organisms, as accurately as possible. This is because reads that align equally well to multiple genomes cannot be classified, but they do indicate that some portion of the sample abundance should be shared among those genomes and not the others.

To ensure that poor alignments are not counted, and that multi-aligned reads have aligned roughly equally-well to all of its potential source organisms, we only consider reads that align at least 95% of bases. We empirically found that this allows slightly more flexibility for errors than requiring perfect alignments, while still retaining high fidelity. As long as one such read aligned uniquely to a genome, we counted that organism as present, while all genomes with no uniquely-aligned reads were discarded. We found this rule to balance precision and recall effectively. We allow users to control how many reads must uniquely align to an organism and what percentage of their bases need to align to count an organism as present, allowing users to modulate false positive rate and false negative rate to suit their needs.

## Authors’ contributions

NL wrote the manuscript and code and carried out the experiments. MA contributed to the software development and manuscript preparation. DK developed CMash and the variant of it used in this method. SM, DK, and NL developed the Metalign method and designed the experiments. NL, SM, DK and EE collaborated on the direction and goals of the project.

## Acknowledgements

The authors would like to thank Dr. Lana S. Martin for her assistance with manuscript and figure preparation.

## Supplementary Information

### Data Availability

We exclusively used publicly-available datasets in this paper. For the CAMI 1 results, we used the results and datasets from the GigaDB dataset here: http://dx.doi.org/10.5524/100344. We used the challenge datasets, not the toy datasets. For the medium complexity datasets, we used the insert size 270 reads. The Peabody mock community data was retrieved using the link in their paper ^1^. The prokaryote-isolated Tara Oceans reads used in this study are available on EBI here: https://www.ebi.ac.uk/ena/data/view/PRJEB1787. The run accessions we used, which were not chosen for any particular reason, were ERR598948, ERR598949, ERR598950, ERR598952, ERR598957. Specifically, we went to the first page in which the Instrument Model for all samples was Illumina HiSeq 2000. We then downloaded all ten datasets on that page, and decided to use only the deep chlorophyll maximum samples, so that the samples would be from a consistent ocean layer. Extra details on each file (sampling location, depth, isolation method, etc) can be found by clicking on the sample accession corresponding to each of those run accessions. Information on the locations of some of the Tara Oceans stations can be found here: http://ocean-microbiome.embl.de/companion.html.

### Replication

We used the docker tag metalign:0.6.2 to generate all results in this paper. We have tried to include all commands run to generate results, raw results (including timing), and intermediate processed results in our GitHub repository, as well as a Jupyter notebook with the code to generate the figures. Because the mock community had very low diversity and reconciling the different output formats of the various methods is difficult, the results for that dataset were compiled via careful manual inspection. However, because the results for the programs are available on GitHub, independent manual or automated validation is possible. For the other methods, we used their default settings (except for number of threads, which we set to 4 for timing purposes on the Tara Oceans data) and their default databases as of October 2019.

### Performance Metrics

We evaluate the performance of multiple methods using several different metrics, which are designed to encompass both performance in the binary classification task of predicting species presence or absence, and in the estimation of relative abundances of taxa. For species presence and absence, a “True Positive” (TP) indicates that a species that is actually present in a sample is correctly predicted as being present by a method, while a “False Positive” (FP) indicates that the method predicted the presence of a species that is absent from a sample and a “False Negative” (FN) indicates that a species was actually present in a sample but a method did not predict its presence. We use two metrics to assess the performance of a method in species presence/absence, precision and recall, defined below. Additionally, we report the F1-Score, which is defined as the harmonic mean of precision and recall. All three of these metrics range from 0 to 1, or 0% to 100%.

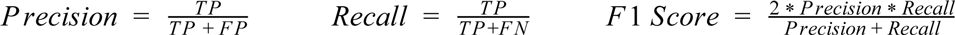

Another common metric is the Jaccard index, which in this case refers to the number of true positives (intersect between predicted and actual communities) divided by the true positives plus the false positives and negatives:

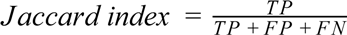

We use the L1-error as a measure of how accurately a method computes the relative abundances of species in a sample. The L1-error is the sum of absolute value differences between predicted species abundances and actual species abundances, and ranges from 0 (completely correct) to 2 (completely incorrect). An L1-error of 0 indicates that the exact set of actual species and their actual abundances is predicted perfectly, while a score of 2 indicates that the set of predicted species is completely incorrect. The L1-error can be described mathematically as:

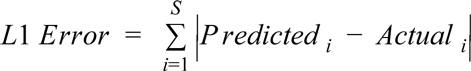

where S is the set of taxa that are predicted or actually present and i is the summation index. Finally, UniFrac ^2, 3^ is a widely-used metric that measures differences between two microbial communities as the fraction of branch lengths of a phylogenetic tree that are present in one community or the other, but not both. More details on UniFrac and Weighted UniFrac can be found in the original papers by Lozupone et al. ^2, 3^.

Krona ^4^ is a popular bioinformatics visualization tool that generates hierarchical charts displaying taxonomic clades and their relative abundances. We used this tool to visualize the results of the methods run on the Tara Oceans data (**Figures S4-S7**), since results cannot be compared due to the lack of known ground truth.

## Supplementary figures

**Figure S1.**
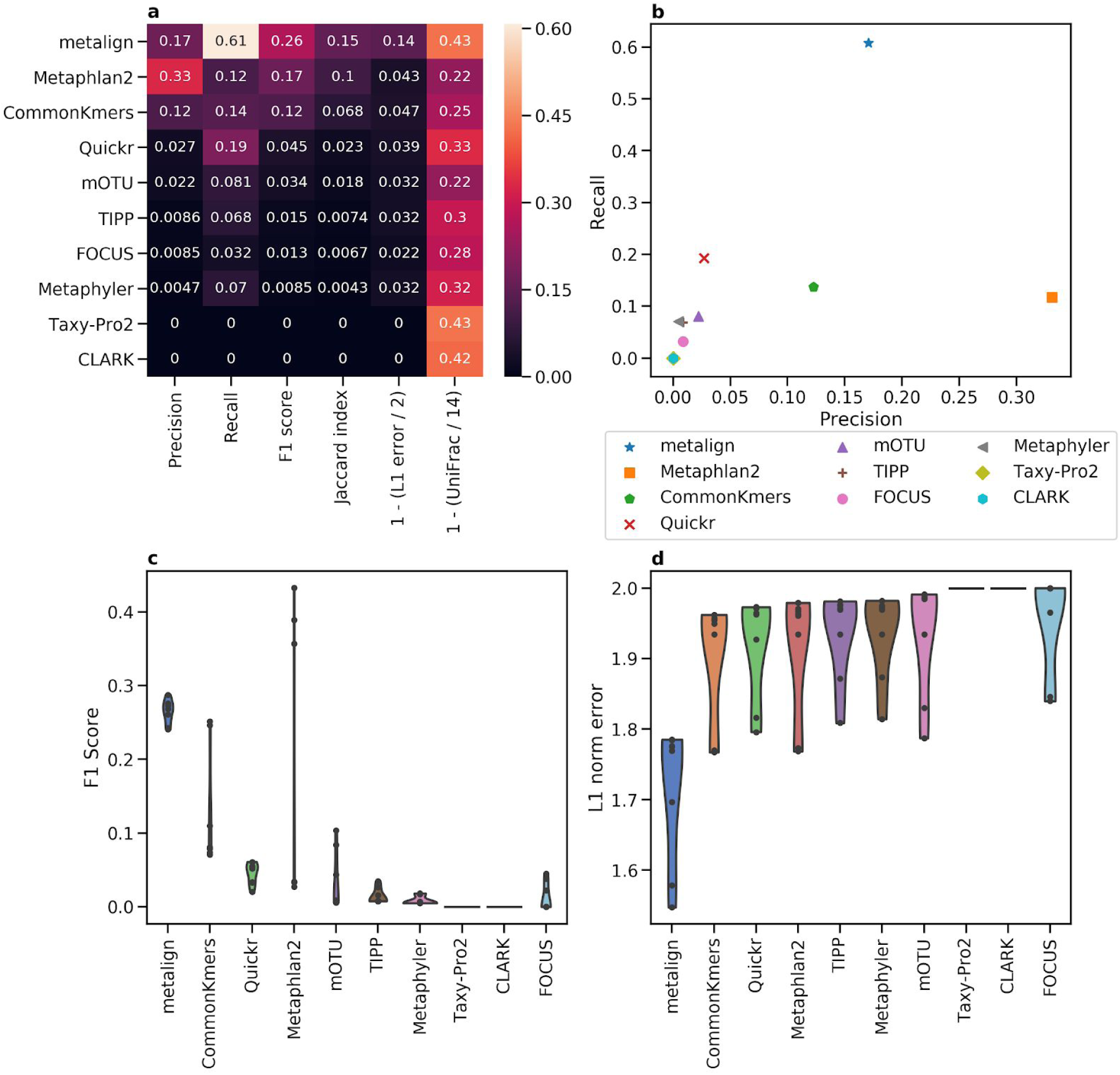
Comparison between Metalign and methods submitted to the first CAMI competition on the species level. (A) Heatmap showing Precision, Recall, F1 score, Jaccard index, and 0-to-1 scaled versions of L1 error and Weighted UniFrac, such that higher is better for each column, averaged across all datasets. (B) A scatter plot of Precision (x-axis) versus Recall (y-axis), averaged across all datasets. (C and D) Violin plots of F1 score (C; higher is better) and L1 error (D; lower is better), respectively, where the x-axis contains the methods, the y-axis represents the metric, and each dot represents one dataset.

**Figure S2.**
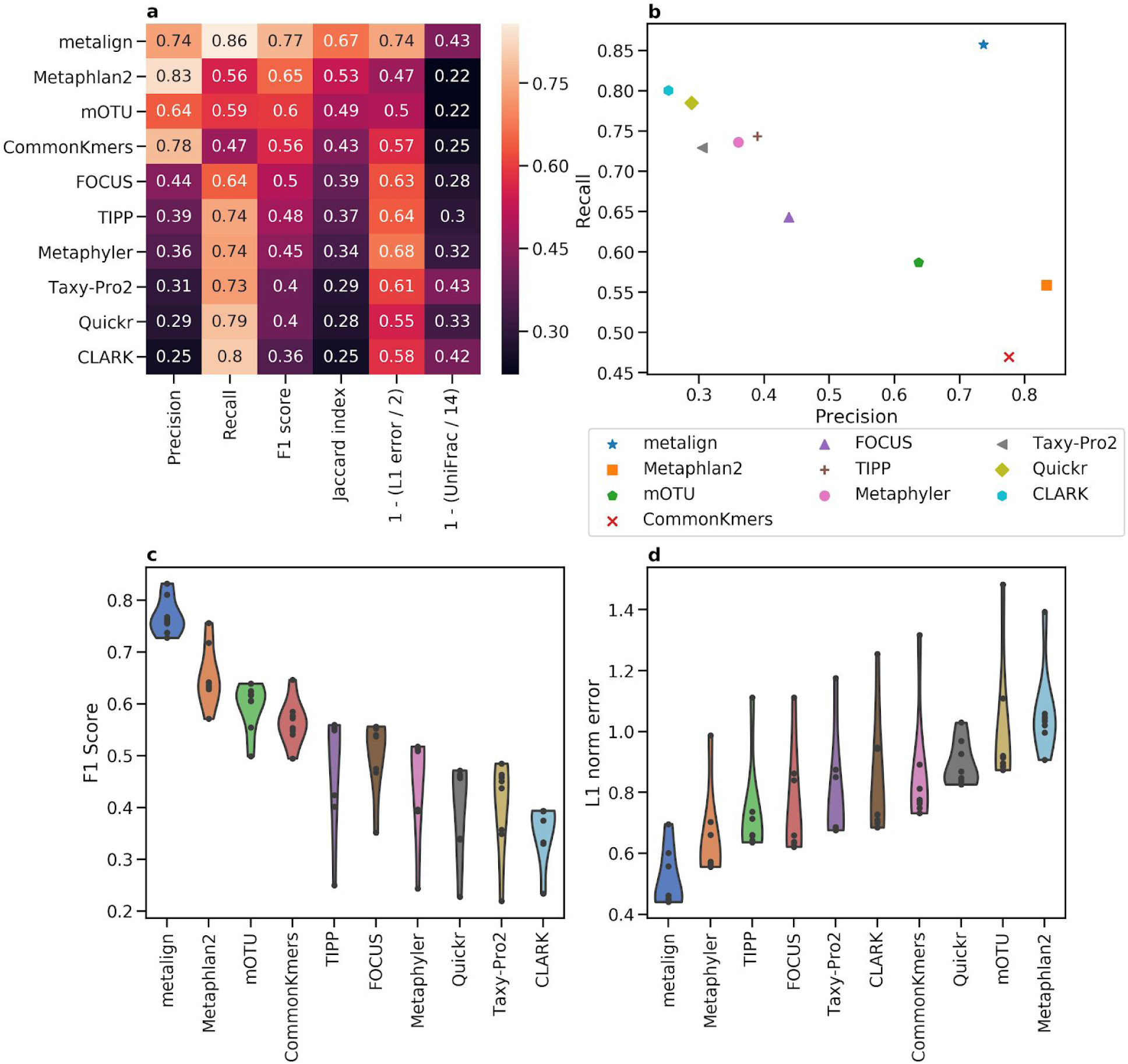
Comparison between Metalign and methods submitted to the first CAMI competition, averaged over all taxonomic levels. (A) Heatmap showing Precision, Recall, F1 score, Jaccard index, and 0-to-1 scaled versions of L1 error and Weighted UniFrac, such that higher is better for each column, averaged across all datasets. (B) A scatter plot of Precision (x-axis) versus Recall (y-axis), averaged across all datasets. (C and D) Violin plots of F1 score (C; higher is better) and L1 error (D; lower is better), respectively, where the x-axis contains the methods, the y-axis represents the metric, and each dot represents one dataset.

**Figure S3.**
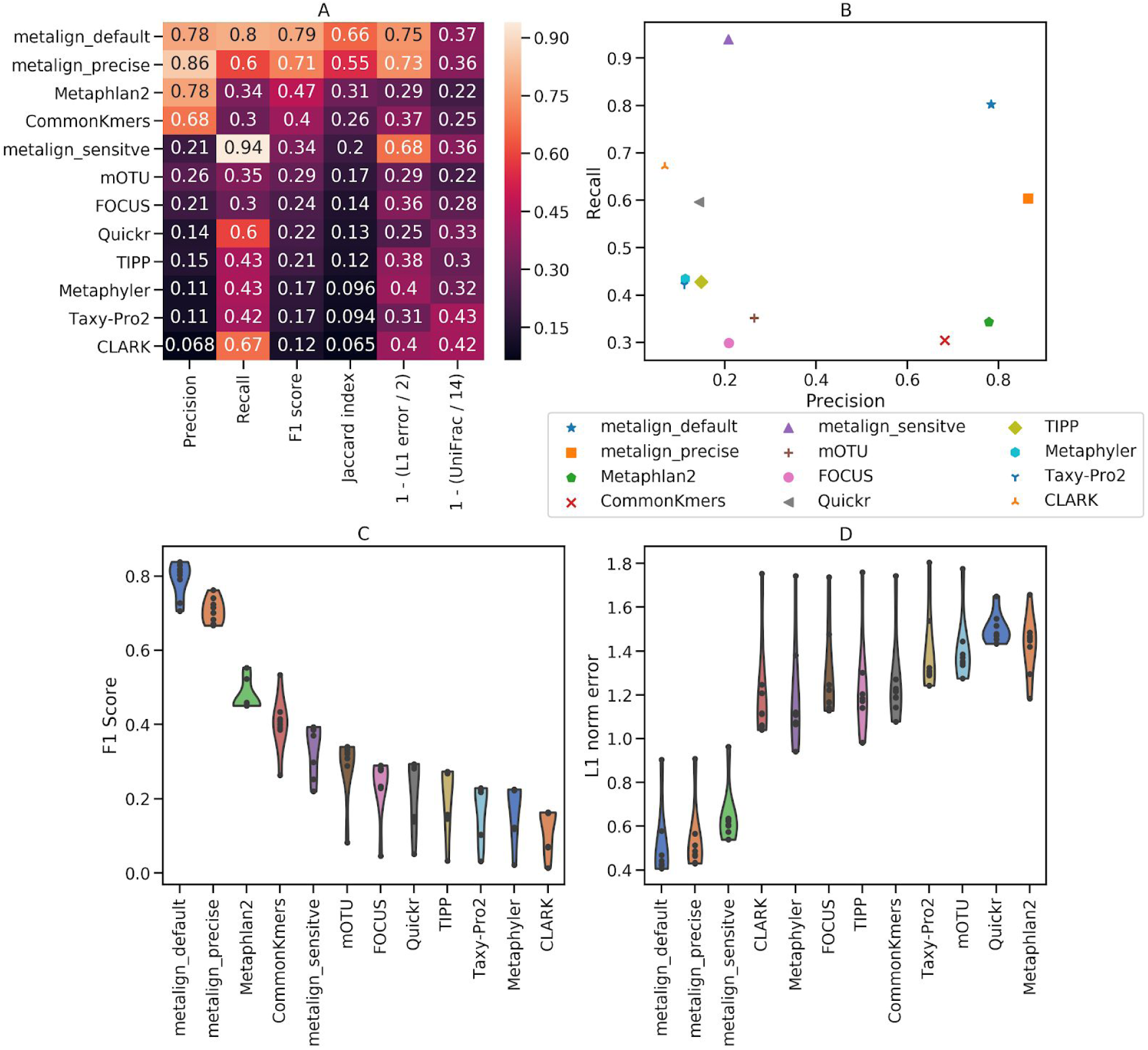
Comparison between Metalign, including its “sensitive” and “precise” settings, and methods submitted to the first CAMI competition, on the genus level. (A) Heatmap showing Precision, Recall, F1 score, Jaccard index, and 0-to-1 scaled versions of L1 error and Weighted UniFrac, such that higher is better for each column, averaged across all datasets. (B) A scatter plot of Precision (x-axis) versus Recall (y-axis), averaged across all datasets. (C and D) Violin plots of F1 score (C; higher is better) and L1 error (D; lower is better), respectively, where the x-axis contains the methods, the y-axis represents the metric, and each dot represents one dataset.

**Figure S4.**
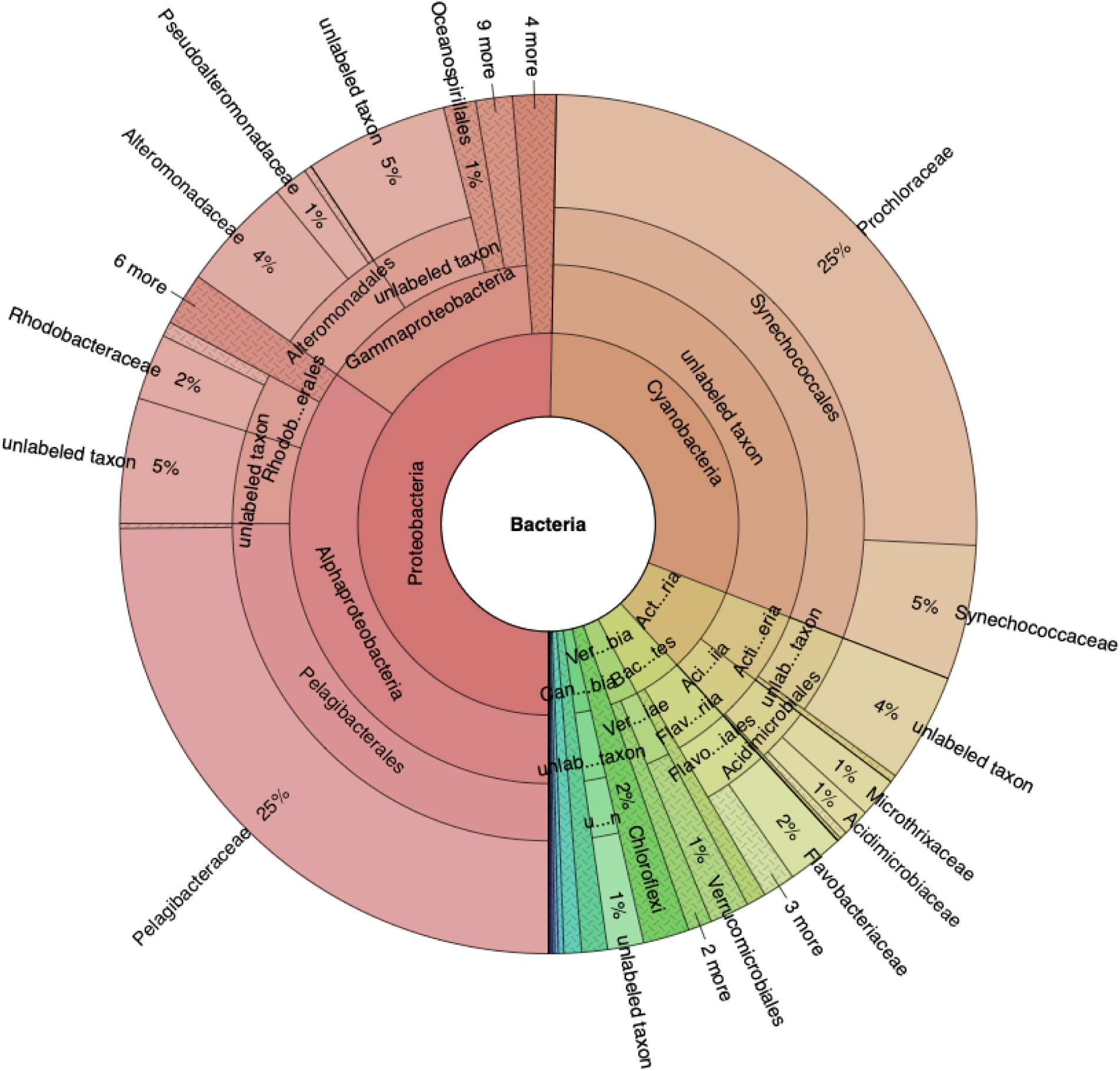
Krona ^4^ plot showing Metalign results on the Tara Oceans ^5^ deep chlorophyll maximum samples. Results shown down to the Family level.

**Figure S5.**
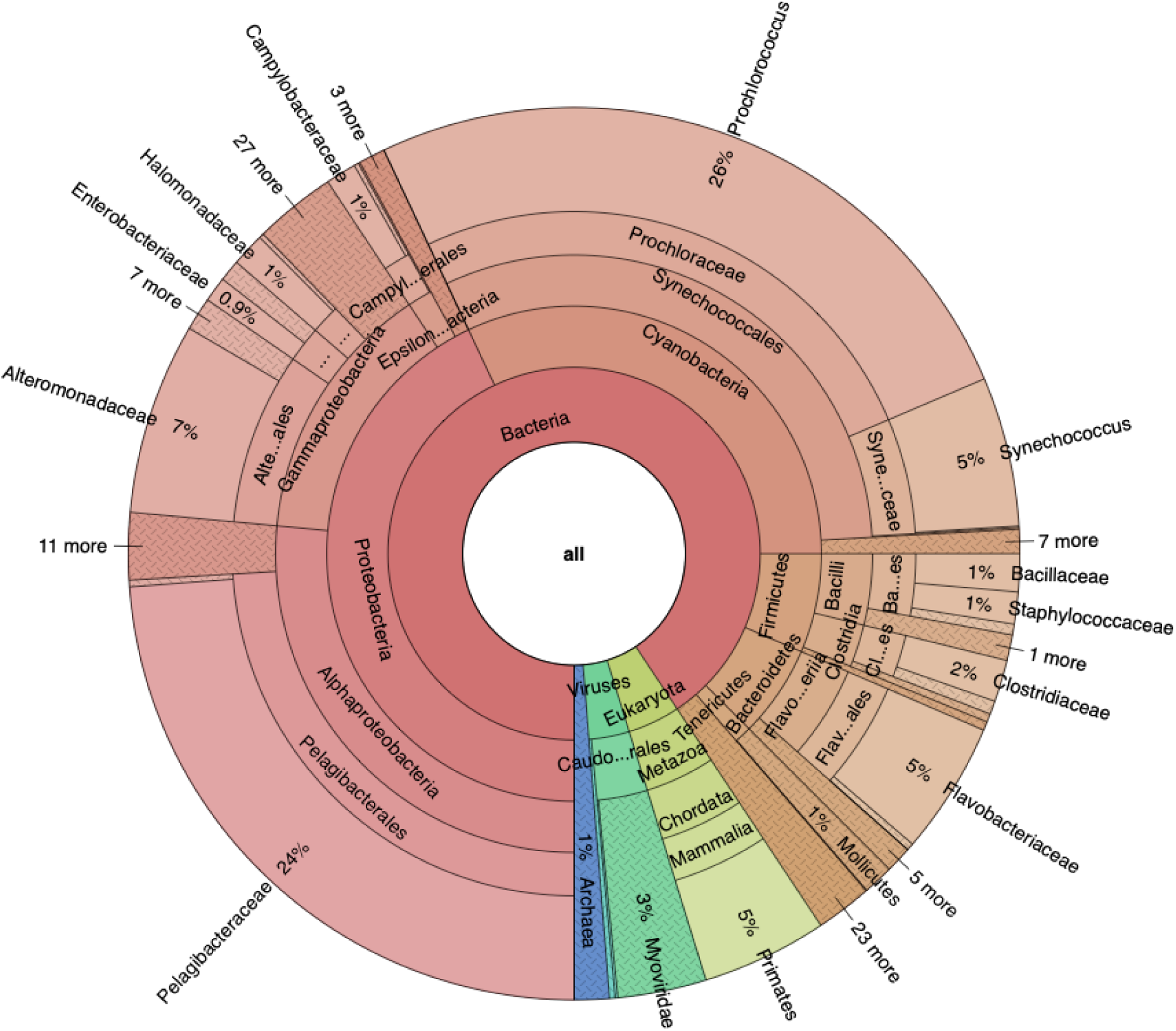
Krona ^4^ plot showing Kraken2 ^6^ + Bracken ^7^ results on the Tara Oceans ^5^ deep chlorophyll maximum samples. Results shown down to the Family level.

**Figure S6.**
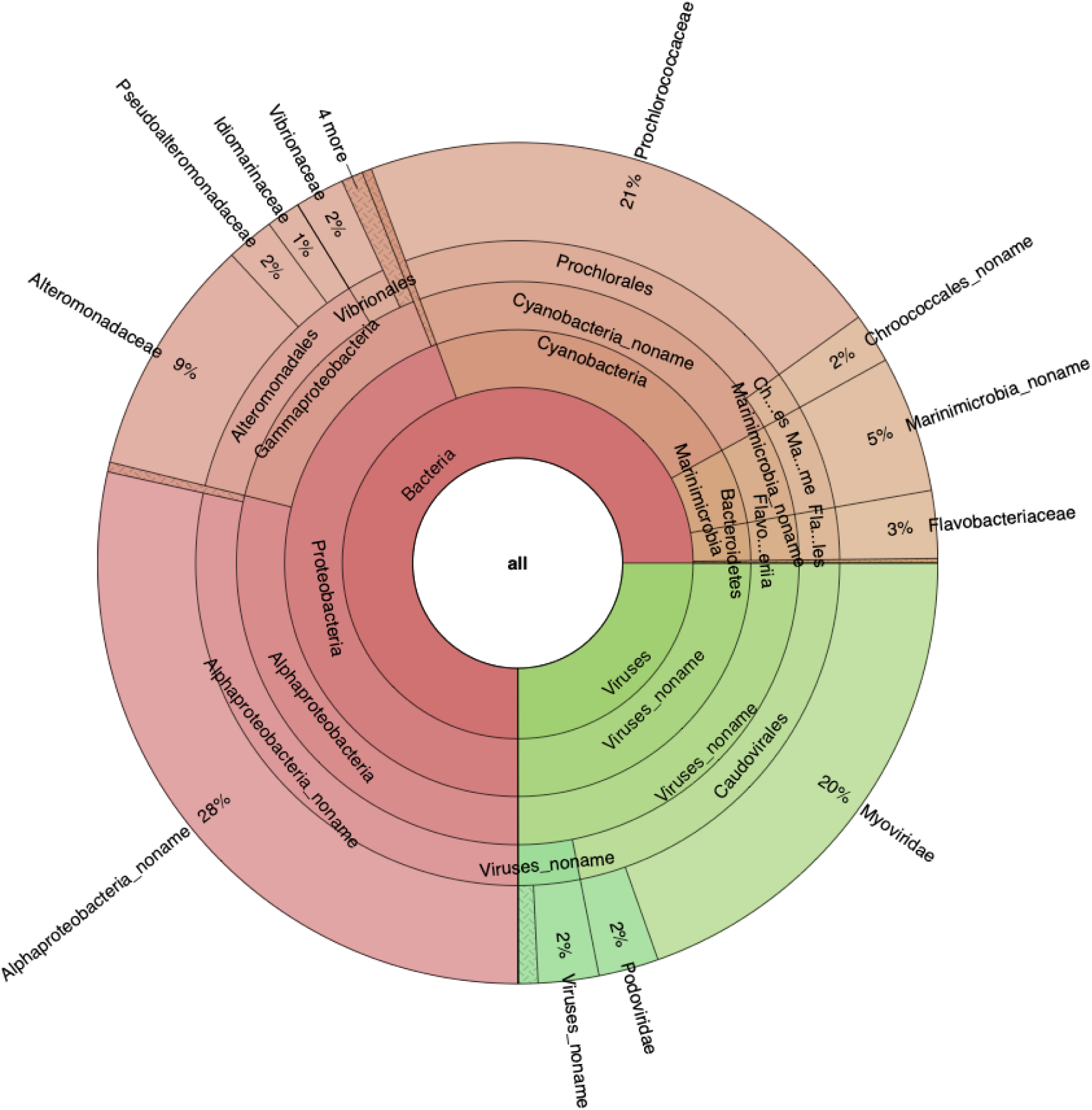
Krona ^4^ plot showing MetaPhlAn2 ^8^ results on the Tara Oceans ^5^ deep chlorophyll maximum samples. Results shown down to the Family level.

**Figure S7.**
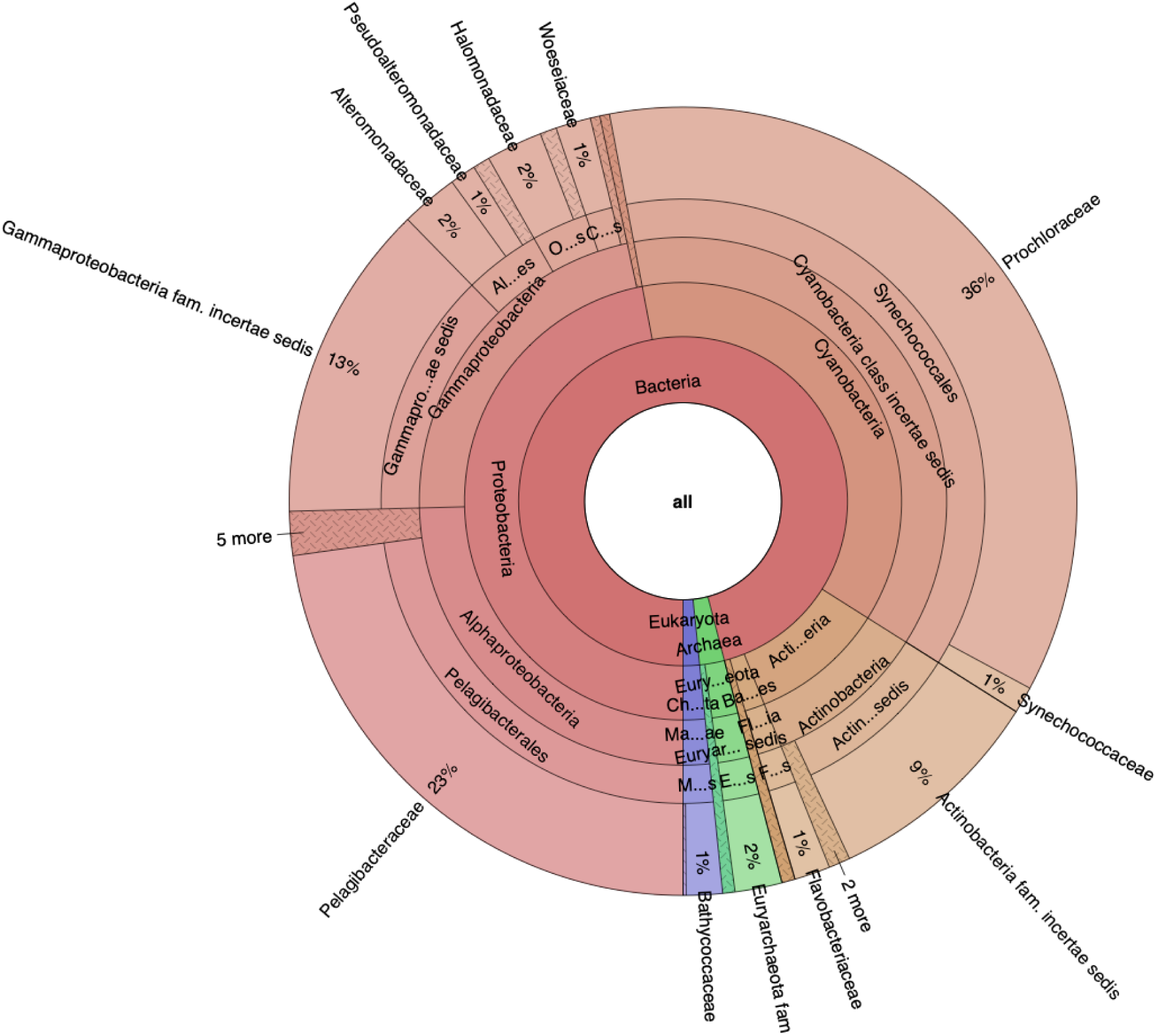
Krona ^4^ plot showing mOTUs2 ^9^ results on the Tara Oceans ^5^ deep chlorophyll maximum samples. Results shown down to the Family level.

